# Modified Elek test improves *in-vitro* detection of diphtheria toxin

**DOI:** 10.64898/2026.02.27.708555

**Authors:** Edgar Badell, Sylvie Bremont, Marion Barbet, Virginie Passet, Chiara Crestani, Sylvain Brisse

**Affiliations:** Institut Pasteur, Université Paris Cité, Biodiversity and Epidemiology of Bacterial Pathogens, Paris, France; National Reference Center for Corynebacteria of the diphtheriae complex, Institut Pasteur, Paris, France; European Reference Laboratory of Diphtheria and Pertussis, Institut Pasteur, Paris, France

**Author notes:** Correspondence: Prof Sylvain Brisse, Institut Pasteur, 25-28 rue du Docteur Roux, 75724 Paris Cedex 15, France.

**Keywords:** Elek’s test, *Corynebacterium ulcerans*, *Corynebacterium diphtheriae*, *Corynebacterium ramonii*, Diphtheria toxin detection, Diphtheria, Laboratory diagnosis

## Abstract

**Purpose:** Diphtheria is caused by toxigenic strains of the *Corynebacterium diphtheriae* complex, mainly *Corynebacterium diphtheriae* and *C. ulcerans*. The diagnosis of diphtheria relies on detecting the diphtheria toxin (DT), for which Engler’s method of Elek’s immunoprecipitation test is the gold standard. A recent optimization of Engler’s method was proposed by Melnikov and colleagues, showing higher sensitivity for *C. ulcerans*. The goal of our study was to test and adapt this optimized method, and to re-analyze apparent non-toxigenic *tox* gene bearing (NTTB) isolates from our collection.

**Methods:** We included 48 *C. ulcerans, C. ramonii* and *C. diphtheriae* isolates previously categorized as NTTB but for which no genetic explanation was found for the lack of DT expression. DT production was tested using Melnikov’s method with further modifications made by us: *i*) increasing the antitoxin concentration; *ii*) using 5°C as the incubation temperature after 24h; and *iii*) modifying the layout of control and test strains on agar plates.

**Results:** 35 of 38 *C. ulcerans*, 3 *C. ramonii* and 8 of 10 *C. diphtheriae* were found to be toxigenic. No genetic explanation was found regarding two non-toxigenic isolates (1 *C. diphtheriae* and 1 *C. ulcerans*), whereas for one *C. diphtheriae, IS1132* was detected upstream of the *tox* gene.

**Conclusion:** Our modified implementation of Melnikov’s Elek test improved our ability to detect diphtheria toxin production. Most isolates previously considered as NTTB but with no genetic explanation, were shown to be toxigenic using the novel method.

## Introduction

Diphtheria is caused by toxigenic strains of several species of the *Corynebacterium diphtheriae* species complex, mainly *Corynebacterium diphtheriae* and the emerging zoonotic pathogen *C. ulcerans*. Diphtheria toxin (DT), the major virulence factor of these bacteria, is responsible for systemic diphtheria symptoms, such as cardiac, renal and neurological ones. Toxigenic strains harbor a *tox* gene-carrying bacteriophage and produce DT; however, some *tox* gene carrying isolates do not produce the toxin and are called non-toxigenic *tox* gene bearing (NTTB) [1,2]. Detection of DT is central to diphtheria diagnosis [3] and defines patient management and public health measures. It is therefore critical to identify toxigenic *Corynebacterium* strains accurately. Although the diphtheria toxin gene (*tox*) can be detected by PCR, this approach cannot distinguish between toxigenic and NTTB strains [3].

The Elek test (or Elek’s test) was designed in 1949 to detect the production of DT [4]. This immunoprecipitation assay relies on the interaction on agar plates between the DT and antitoxin polyclonal antibodies, which results in the formation of a visible precipitate. The original method was modified into the current gold standard for DT detection by Engler and colleagues in 1997 [5]. In our laboratory, Engler’s protocol has been used since 2008, leading to identifying several NTTB isolates, particularly in *C. ulcerans*, in most cases with no obvious genomic explanation for DT production deficiency.

Recently, Melnikov and colleagues [6] described an optimization of Engler’s method and reported DT production in *C. ulcerans* clinical isolates, which had initially been considered as NTTB.

The goal of this study was to re-analyze 48 apparent NTTB isolates from our collection, which had no apparent disruption of the *tox* gene, with Melnikov’s implementation of Elek’s test. While doing so, we further introduced modifications to the experimental protocol.

## Materials and methods

### Isolates

Thirty-five *C. ulcerans*, 3 *C. ramonii* and 10 *C. diphtheriae* non-duplicate NTTB isolates, mostly from the collection of the French National Reference Center for Corynebacteria of the *diphtheriae* complex were used (**Table 1**). The *C. diphtheriae* and *C. ramonii* were all of human origin whereas the *C. ulcerans* were both of human (n=22) and animal (n=13) origins. Clinical isolates were collected in France between 2008 and 2024, and three reference strains were included from the Collection of Institut Pasteur (CIP), isolated in 1951 or 1994 (**Table 1**). Genomic sequences of 43 isolates were previously obtained [7,8] and available from public sequence repositories, whereas the 5 remaining ones were obtained in this work using the same method as in [7] (**Table 1**).

**Table 1.**
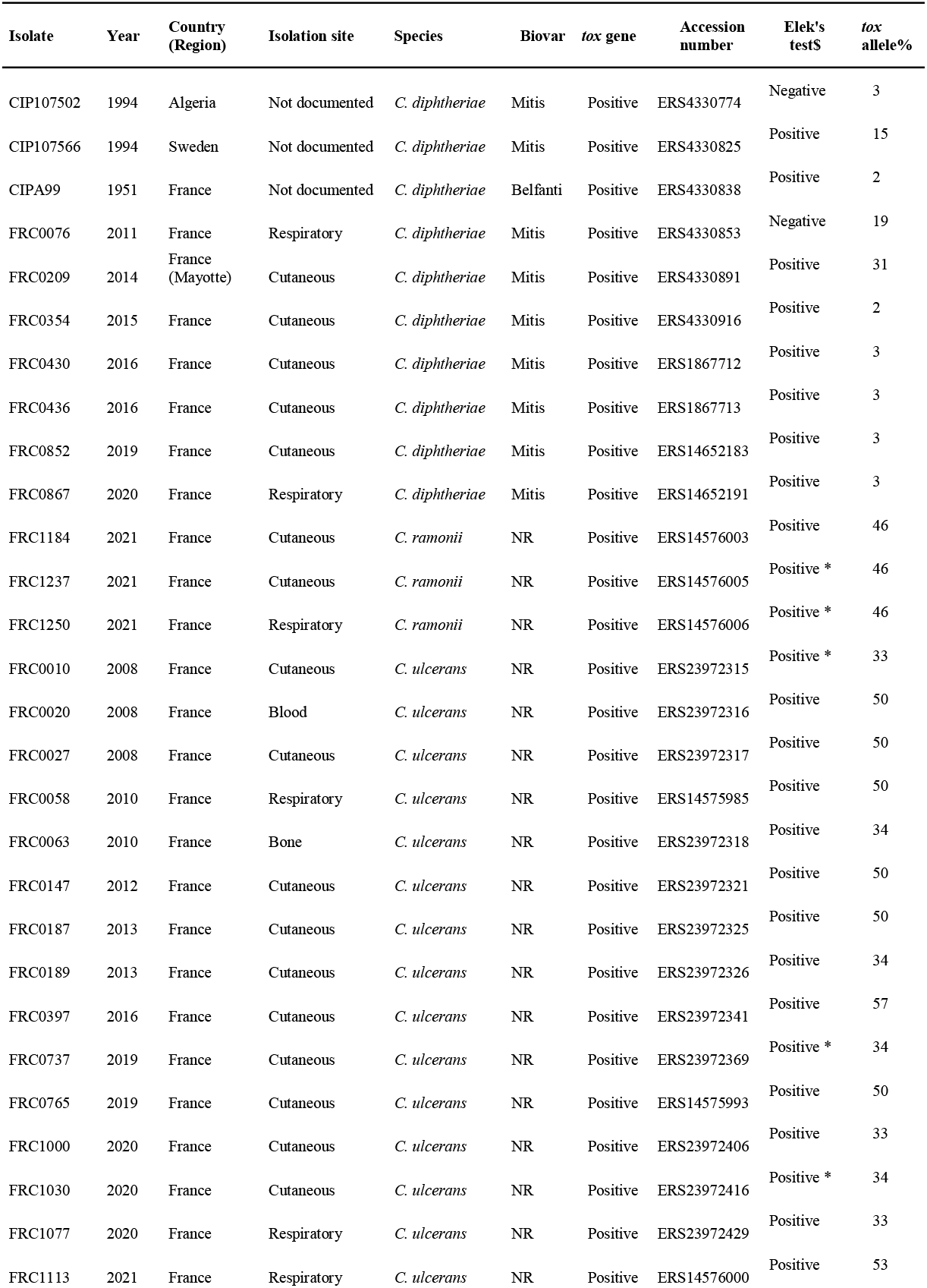
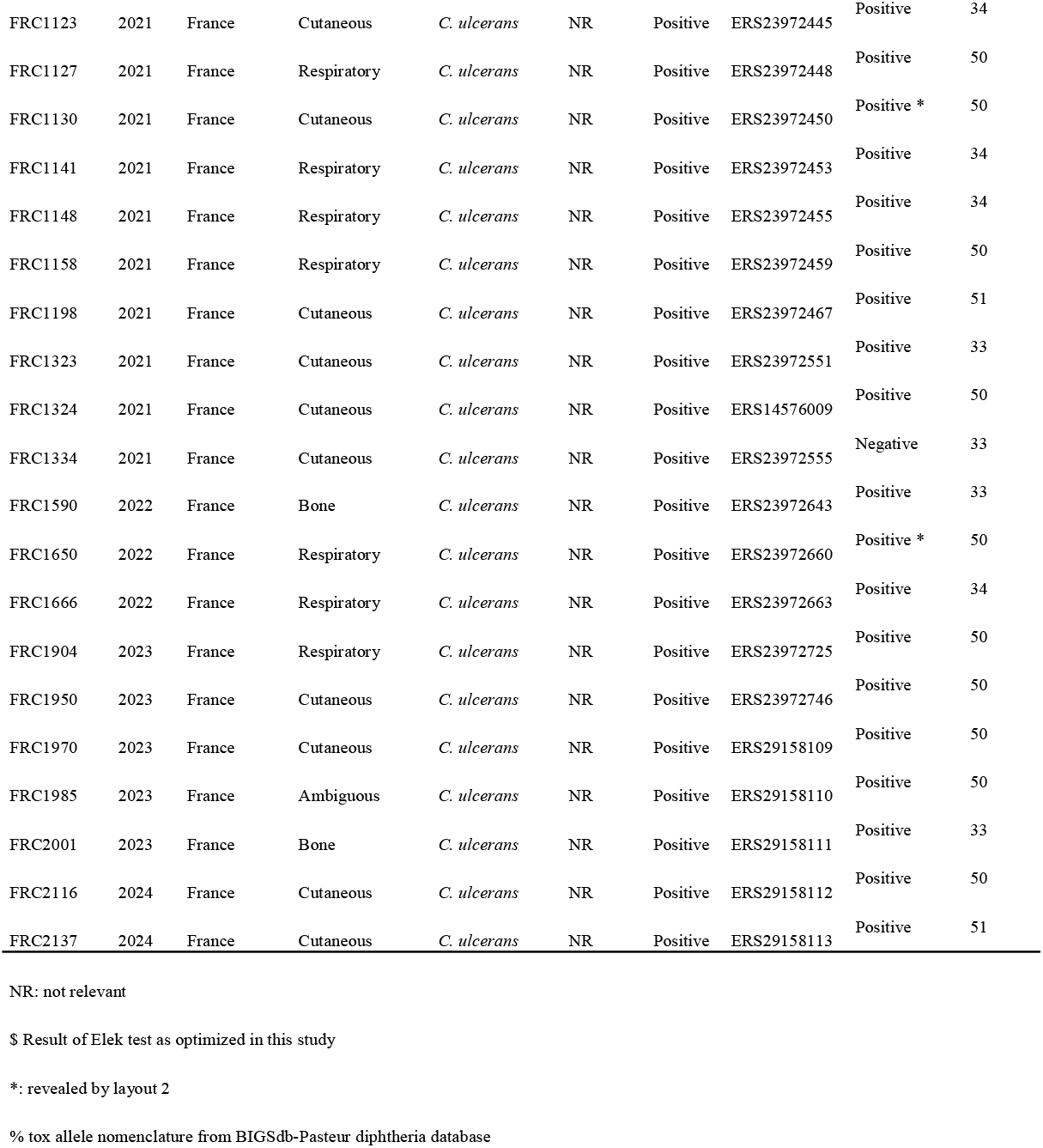
*Corynebacterium* strains used in this study and their characteristics.

### *tox* gene presence and Elek tests

The presence of the *tox* gene was determined by PCR [9] and confirmed after genomic sequencing using the bioinformatics tool diphtOscan [10]. DT production was prospectively determined using Engler’s method [5]. In this study, we reanalyzed DT production using a modified version of Melnikov’s optimized Elek test [6], which we describe below.

Bacteria were grown on Columbia blood agar (bioMérieux) and Tryptic Soy Agar (in-house-made medium) for 24 h at 35°C. The Elek agar base was prepared as described previously [5]. Briefly, a molten Elek agar (4 ml) cooled to 50°C was supplemented with 1 ml (20% final concentration) of newborn bovine serum (MP Biomedicals Europe, France), mixed gently and poured into 5.5 cm diameter Petri dish (Corning Gosselin®, France). The medium was spread over the bottom by gently swirling the plate. The plate was then left in a laminar flow hood for one hour to solidify the agar medium, with the plate lid semi-open to dry the agar surface and avoid condensation. Such plates can be stored for 5 to 8 days at 4°C.

Paper discs of 6 mm (Mast group Ltd, UK) were soaked with a diphtheria antitoxin serum (DAT) from Vins Bioproducts Limited, India. Impregnated discs were dried in a laminar flow hood for about 2 h. Once dried, discs can be stored for 6 months, between 2°C and 8°C, in a sealed container.

Two concentrations of DAT were tested: (i) 2.5 UI, as recommended in Melnikov’s 2022 version of Elek’s test; this concentration was achieved by depositing on each paper disc, 25 μL of DAT at a concentration of 100 UI/ml in sterile distillate water; and (ii) 12.5 UI per disc (500 UI/ml).

Elek’s test was initially performed as described by Melnikov and colleagues [6] on a few test strains. Briefly, a paper disc containing the DAT was placed on the agar surface and bacteria were heavily inoculated on 6 spots around the antitoxin disc, using disposable plastic loops with a capacity of 1 μl, following the layout proposed by Melnikov and colleagues (**Figure 1, left**). In this layout, 3 of the 6 spots contained the toxigenic control strain, 1 contained the non-toxigenic control strain, and 2 spots contained the test isolate. Both the inoculum to antitoxin disc distance, and the diameter of the bacterial inoculation spots, were set at 6 mm.

**Figure 1.**
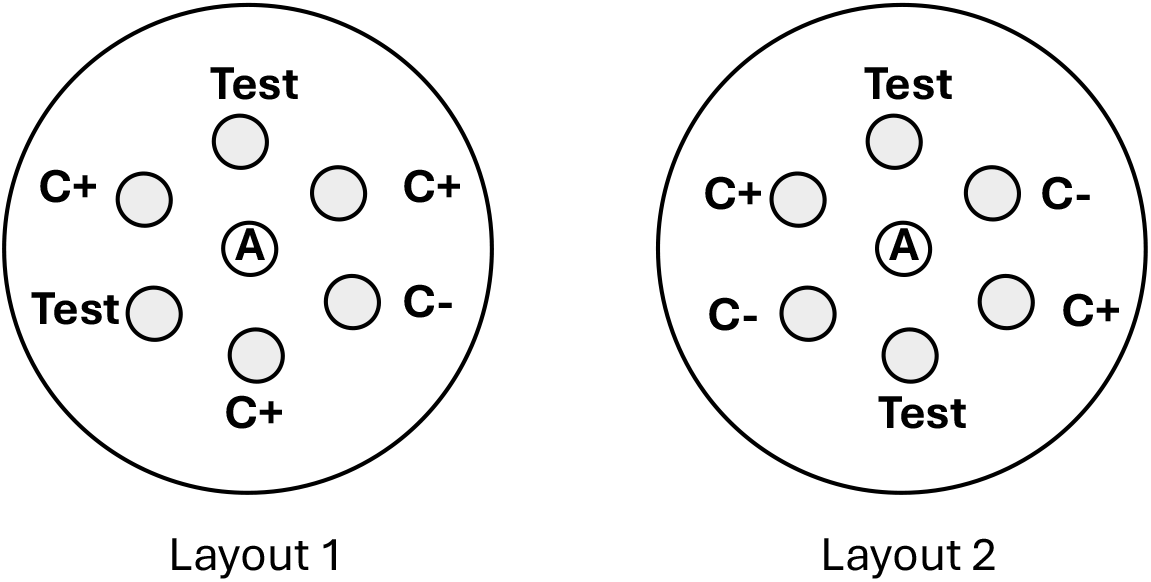
Layouts used for Elek’s test. Left: layout 1 used by Melnikov *et al*. 2022. Right: layout 2 proposed herein. Test: isolate tested; A: anti-toxin impregnated disc; C+: positive control strain; C-: negative control strain.

Because in some tests, bacterial growth overlapped the precipitin lines, preventing its observation, we modified Melnikov’s implementation by reducing the incubation temperature to 5°C beyond the first 24 hours (instead of 35°C).

Reference strains NCTC 10648 (toxigenic *C. diphtheriae*, biovar Gravis) and NCTC 10356 (*tox* gene-negative *C. belfantii*) were used as positive and negative controls, respectively, for all versions of Elek’s tests. All experiments were carried out in duplicate.

Elek’s test was considered positive if a precipitin line was visible between the bacterial spots and the paper disk loaded with antitoxin. The precipitin line should merge with the lines formed by the positive control colonies located on both sides of the test isolate after 16 to 24h of incubation at 35°C; or, if the precipitin line was not well defined at 24 h, the incubation was extended to 48 h. Further, if after 48 h of incubation the precipitation lines were still not clearly visible, the strain was retested after changing the position of control strains on agar plates as shown in the right panel of Figure 1.

## Results

We first tested the method of Melnikov and colleagues [6], using 5 *C. ulcerans* NTTB strains. In our hands, when using discs containing 2.5 UI of DAT, the immunoprecipitation lines for positive controls as well as for the 5 tested isolates were ambiguous even after 48h of incubation. We thus increased the quantity of DAT to 12.5 IU per disc, which resulted in clearly visible immunoprecipitation lines. Therefore, the quantity of 12.5 IU was used in subsequent experiments.

The precipitin lines were typically produced after 16–18 h at 35°C for the toxigenic control as well as for 31 isolates (**Figure 2**). For 6 other isolates, non-ambiguous precipitin appeared after 24 h of additional incubation at 5°C. However, for 5 *C. ulcerans* and *2 C. ramonii* isolates, precipitin lines were still not clearly visible even after this extended incubation at 5°C, whereas for 3 isolates (*C. ulcerans* FRC1334, and *C. diphtheriae* CIP107502 and FRC0076) the result was considered negative. No genetic explanation was found for CIP107502 and for FRC1334, whereas for FRC0076, insertion sequence IS1132 was detected right upstream of the *tox* gene. This insertion could explain the Elek-negative result due to the disruption of the *tox* promoter region.

**Figure 2.**
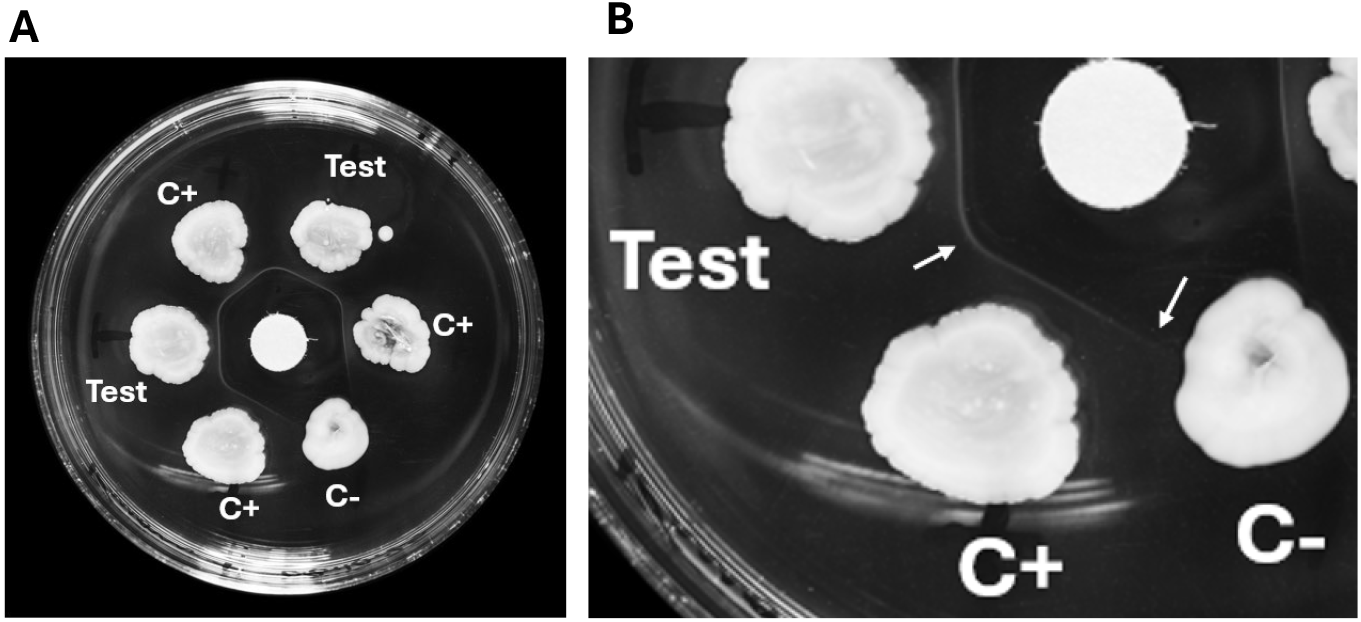
Results of Elek’s test using layout 1. **A.** Results obtained with an isolate that produces high levels of DT. A continuous precipitation line was formed in front of the tested isolate and the control strain (C+). **B.** Magnification of (A). The arrows point to the differences observed between the edges of the precipitation lines: the precipitation line is curved between the tested isolate and C+, whereas it is straight between C+ and C-.

We next retested the 10 latter isolates using a novel positioning layout (**Figure 1, layout 2**). The observed incurvation of the precipitin lines suggested that the 5 *C. ulcerans* and *2 C. ramonii* isolates with ambiguous results with layout 1, were toxigenic (**Figure 3**).

**Figure 3.**
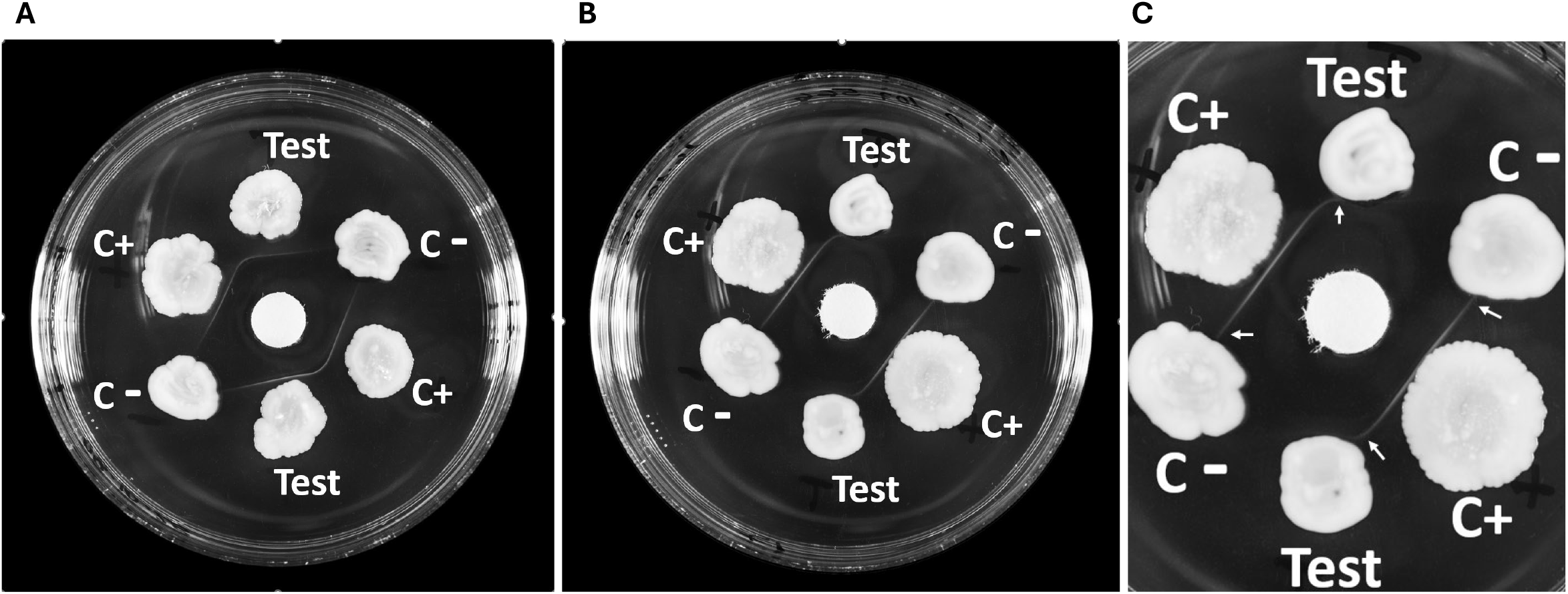
Results of Elek’s test obtained using layout 2. **A.** Easily visible precipitin lines obtained with isolate FRC1148 **B.** Results obtained with isolate FRC1030, for which the precipitin line is not apparent **C.** Magnification of (B). Arrows point to the edges of the precipitin lines, which are either curved (between C+ and the test isolate), or straight (between the test isolate and C-). This observation suggests that the test isolate produces DT at low levels.

The modifications introduced in our approach can be summarized as follows: we used 12.5 UI of DAT deposited on each paper disk, an incubation at 5°C between 24h and 48h timepoints, and a novel layout that disambiguates the weakly positive results.

Overall, when considering the 48 tested isolates, 34 out of 35 *C. ulcerans*, 8 out 10 *C. diphtheriae* and 3 *C. ramonii* were determined as being toxigenic.

## Discussion

The goal of this study was to evaluate in our laboratory, Elek’s test protocol recently proposed by Melnikov and colleagues [6] to improve detection of DT production. Further, we aimed to reevaluate apparent NTTB isolates from our collection. Preliminary experiments with five test isolates showed that in our hands, using Melnikov’s protocol, no easily interpretable results were obtained.

Among the 3 modifications we introduced, one consisted of increasing DAT concentration from 2.5 UI to 12.5 IU per disc. The need to increase the quantity of DAT per disc to 12.5 UI might be due to different batches of antitoxin used here and in Melnikov’s study; quantification of antitoxin levels in DAT products is not performed in our laboratory, and we had to rely on the provider’s quantification information. Note that we used an enzyme-refined, purified DAT different from the one used by Melnikov and colleagues, the latter one being not available commercially.

A second modification we introduced involved reducing the incubation temperature from 35°C to 5°C between 24 h and 48 h. A low temperature slows down the bacterial growth and prevents bacteria from overlapping with the precipitation lines, which hindered the observation of their edges in our initial experiments.

The two modifications described above allowed obtaining clear positive results of DT production for 28 of 35 *C. ulcerans* and 1 *C. ramonii* isolates, which had previously been considered as NTTB. However, 7 isolates (5 *C. ulcerans* and *2 C. ramonii*) still showed very diffuse immunoprecipitation lines that were difficult to observe. We nevertheless noted that the immunoprecipitation lines of the positive control continued to extend toward each of these 7 isolates, taking a curve, in the same manner as when the positive control was placed besides a clearly toxigenic strain. This incurvation was not observed between the positive and negative controls. Thus, the incurvation suggested that the 7 isolates produce DT at very low levels. To reveal this phenomenon more clearly, we used a new layout, which facilitates the observation of contrasting shapes of the edges of the immunoprecipitation lines formed by the positive control toward the test isolate *versus* a negative control (**Figure 3**). We thus propose, in the absence of *tox* mutations suspected to lead to DT deficiency, the following interpretation: if the edge of the line of the positive control close to the test isolate becomes curved towards the isolate, this isolate can be considered toxigenic even though its own precipitation line is not observed. On the contrary, if the edge remains straight, it can be concluded that the test isolate does not produce DT. Based on this approach, the 7 retested *C. ulcerans* and *C. ramonii* isolates were considered positive for DT production.

The detection of weakly DT-producing isolates is important, as their potential pathogenic role should not be underestimated. DT is lethal for susceptible animals as well as unvaccinated humans at doses of 100 ng/kg or less [11]. Besides, a weak DT production *in vitro* test does not exclude the possibility that the isolate would produce large amounts of DT *in vivo*.

Melnikov and colleagues tested NTTB *C. ulcerans* strains, but not NTTB *C. diphtheriae* or *C. ramonii* strains. Here, we showed that improvements of Elek’s test also allowed detecting DT production in eight *C. diphtheriae* and three *C. ramonii* previously considered as NTTB.

Regarding the two non-toxigenic *C. diphtheriae* isolates, one (FRC0076) presented IS1132 inserted just upstream the complete *tox* gene (allele *tox*-19). In the BIGSdb-Pasteur diphtheria database, the *tox*-19 allele was also found in 2 isolates from the USA, 1 isolate from Spain and one from Germany, all presenting IS1132 upstream of the *tox* gene. The 2 strains from the USA were also described as negative for Elek’s test. No information was available for the Spanish strain, whereas the German isolate was described as Elek-positive.

The second negative *C. diphtheriae* strain, CIP107502, and the *C. ulcerans* isolate FRC1334 had no disruptive mutations in their *tox* gene or in their promoter region. The *tox* alleles of these isolates are *tox*-3 and *tox*-33, respectively. Both alleles are present in dozens of other *C. diphtheriae* or *C. ulcerans* isolates of our collection, most of which are toxigenic. The *dtxR* gene, coding for the master regulator DtxR, is also typical in these isolates. An explanation for the negative DT detection result in these two isolates might lie in another part of the genome.

The nonspecific precipitation lines observed in Melnikov’s study with unpurified DAT could be due to the diffusion of other bacterial components recognized by the polyclonal antitoxin [6]. The new layout we have introduced here may also help distinguish between specific and nonspecific precipitation lines. Indeed, if the precipitin lines are non-specific, the edges of the line will be curved between the negative control spot and its adjacent strains, whereas specific lines edges will be straight [6]. Here, we performed a total of 142 tests using a unique batch of anti-toxin and did not observe nonspecific precipitation lines, neither in the tests described above (48 isolates in duplicate), nor for 22 additional *tox*-negative isolates: 2 *C. belfantii*, 9 *C. diphtheriae* biovar Gravis, 5 *C. diphtheriae* biovar Mitis, 1 *C. pseudotuberculosis*, 2 *C. ramonii*, 1 *C. rouxii* and 2 *C. ulcerans* (*data not shown*). However, when using another batch of DAT subsequently, we observed the presence of nonspecific precipitation lines with some strains when incubation was prolonged to 48 hours (data not shown). Therefore, the occurrence of nonspecific precipitation lines may be antitoxin batch dependent.

In conclusion, we have built on Melnikov *et al*. method [6] and modified (i) the quantity of antitoxin; (ii) the incubation temperature after 24h; and (iii) the position of control and test strains on Elek agar plates. Using these modifications, we detected DT production for most *tox*-positive isolates with no mutation in the *tox* gene, including for *C. ulcerans* and *C. ramonii*, for which toxin production is more difficult to detect. We suggest these modifications may enable more accurate diagnosis and better surveillance of diphtheria. Our results also suggest the existence of rare but so far unknown genetic mechanisms of lack of DT production by *tox*-positive *C. diphtheriae* and *C. ulcerans* isolates.

## Acknowledgements

We thank all previous and current members of the NRC for their contributions to the collection of isolates and their initial characterization, and the unit Image of Institut Pasteur, Paris, for assistance with Elek’s tests photos.

## Funding

The National Reference Center for Corynebacteria of the *diphtheriae* complex receives support from Institut Pasteur and Public Health France (Santé publique France, Saint Maurice, France). The European Reference Laboratory for Diphtheria and Pertussis (EURL-PH-DIPE) receives support from the European Commission through the EU4HEALTH programme (101194675).

## Author contributions

EB and SBri designed the study. EB, SBre and MB conducted the experiments. EB, SBre and MB evaluated the data. VP and CC conducted the genomic analyses. EB wrote the first draft of the manuscript, which was revised by SBri. All authors read and approved the final version of the manuscript.

## Open access

This research was funded, in whole or in part, by Institut Pasteur. For the purpose of open access, the authors have applied a CC-BY public copyright license to any Author Manuscript version arising from this submission.

## Notes

### Competing Interest Statement

The authors have declared no competing interest.

